# A selective GSK3β inhibitor, tideglusib, decreases intermittent access and binge ethanol self-administration in C57BL/6J mice

**DOI:** 10.1101/2024.05.13.593949

**Authors:** Sam Gottlieb, Andrew van der Vaart, Annalise Hassan, Douglas Bledsoe, Alanna Morgan, Brennen O’Rourke, Walker D. Rogers, Jennifer T. Wolstenholme, Michael F. Miles

## Abstract

Over 10% of the US population over 12 years old meets criteria for Alcohol Use Disorder (AUD), yet few effective, long-term treatments are currently available. Glycogen synthase kinase 3 beta (GSK3β) has been implicated in ethanol behaviors and poses as a potential therapeutic target in the treatment of AUD. Here we investigate the role of tideglusib, a selective GSK3β inhibitor, in ethanol consumption and other behaviors. We have shown tideglusib decreases ethanol consumption in both a model of daily, progressive ethanol intake (two-bottle choice, intermittent ethanol access) and binge-like drinking behavior (drinking-in-the-dark) without effecting water intake. Further, we have shown tideglusib to have no effect on ethanol pharmacokinetics, taste preference, or anxiety-like behavior, though there was a transient increase in total locomotion following treatment. Additionally, we assessed liver health following treatment via serum levels of alanine aminotransferase, aspartate aminotransferase, and alkaline phosphatase and showed no effect on aminotransferase levels though there was a decrease in alkaline phosphatase. RNA sequencing studies revealed a role of GSK3β inhibition via tideglusib on the canonical Wnt signaling pathway, suggesting tideglusib may carry out its effects on ethanol consumption through effects on β-catenin binding to the transcription factors TCF3 and LEF1. The data presented here further implicate GSK3β in alcohol consumption and support the use of tideglusib as a potential therapeutic in the treatment of AUD.

## INTRODUCTION

More than 10% of the United States population over age 12 are estimated to meet criteria for Alcohol Use Disorder (AUD)(1). AUD is characterized by uncontrolled ethanol consumption leading to negative social and health consequences, with alcohol being the third-leading cause of preventable death in the United States(2). Despite these tolls, few treatments for AUD exist, with no new FDA-approved therapeutic treatments within the last 18 years. Additionally, the limited treatments we do have are modestly effective at best in reducing alcohol consumption and rates of relapse(3).

To elucidate mechanisms underlying the neurobiology of AUD, our laboratory has previously performed genome-wide expression network profiling of brain regions in mice following acute and chronic ethanol exposure. Those studies reveal the serine-threonine kinase glycogen synthase kinase 3-beta (Gsk3β) to be a hub gene in a network highly regulated by acute ethanol in medial prefrontal cortex (mPFC)(4–6). Cross species analysis further supports Gsk3β as a candidate gene contributing to AUD(7), and human genome-wide association studies identified a Gsk3β-centric network associated with risk for alcohol dependence(6).

GSK3β is ubiquitously expressed throughout the body but has particularly high expression within the brain(8). Initially discovered for its actions in decreasing glycogen synthesis(9), GSK3β is now known to be involved in many processes including neuronal differentiation, synapse development, and both short- and long-term synaptic plasticity(10). GSK3β dysregulation has long been implicated in neurological and psychiatric disorders such as bipolar disorder, schizophrenia, major depressive disorder, and Alzheimer’s disease(11). Previous work on substance use disorder revealed cocaine and morphine both activate GSK3β (12, 13), and this activation is required for development of cocaine-induced conditioned place preference and tolerance to morphine’s antinociceptive effects(14, 15).

In regard to alcohol, gene targeting studies in mice further implicate GSK3β as a modulator of ethanol-related behaviors. Viral-mediated overexpression of GSK3β within the mPFC increases ethanol consumption in male mice. Lithium, which pharmacologically inhibits both the alpha and beta paralogs of GSK3, decreases ethanol self-administration in these GSK3β overexpressed mice(6). Additionally, the more selective GSK3β inhibitor TDZD-8 decreases operant self-administration of ethanol in rats(6). These prior findings, together with those showing ethanol-induced regulation of GSK3β expression and activity, strongly implicate GSK3β as a potential target for therapeutic intervention in AUD.

Tideglusib, also known as NP031112 or NP-12, is a highly selective, non-ATP competitive, GSK3β inhibitor. While tideglusib has structurally similarity to TDZD-8, it has greater bioavailability(16, 17). Tideglusib is already undergoing clinical trials for various neurodegenerative and developmental disorders(18–22). Here we report actions of tideglusib on ethanol consumption in both a two-bottle choice, intermittent ethanol access and a drinking in the dark (DID) model and investigate tideglusib-induced responses in other behaviors to further assess its potential as a therapeutic agent for AUD. We additionally investigate tideglusib modulation of downstream GSK3β targets involved in synaptic plasticity and neurotransmission. Our findings further support GSK3β inhibitors such as tideglusib as candidate therapeutics in AUD.

## MATERIALS AND METHODS

### Animals

Rodent animal studies and procedures were approved by the Institutional Animal Care and Use Committee of Virginia Commonwealth University and followed the NIH Guide for the Care and Use of Laboratory Animals. Male and female C57BL/6J mice, purchased from Jackson Laboratories (Bar Harbor, ME) were used for all studies, with the exception of intermittent ethanol access (IEA) experiment two and blood ethanol content (BEC) studies which utilized only males. Mice were housed in a temperature and humidity-controlled room in accordance with the Association for Assessment and Accreditation of Laboratory Animal Care approved animal care facility and had ad libitum access to food and water. Animals were housed on utilized Tekland Laboratory Grade Sani-Chips bedding. Animals were habituated to the vivarium at least 7 days prior to beginning experimental manipulations. Mice were housed in groups of 4 for all experiments with the exception of ethanol-drinking studies and taste preference where they were single housed. Mice in drinking studies were kept under a 12-hour reverse light/dark cycle (lights off at 7:00 am) schedule, with mice undergoing other behavioral tests kept under a normal light/dark cycle (lights on at 7:00am). Mice were 8 to 10 weeks of age and weighed approximately 20-30 g at the start of each experiment. All experiments were performed between 7:00 am and 7:00 pm.

### Drugs

Ethanol (EtOH) for drinking studies was prepared as a 20% (v/v) solution in tap water and provided to animals for voluntary oral consumption. For other studies utilizing EtOH, it was made up in 0.9% saline and delivered via intraperitoneal (i.p.) injection. The GSK3β inhibitor tideglusib (Selleck Chemicals, Houston, TX) was prepared for gavage by suspension at 20mg/ml in 26% peg-400 (Sigma), 15% Cremophor EL (Sigma), and water for IEA experiment one and blood ethanol content studies. Tideglusib was prepared in corn oil for all other studies at 20mg/ml with the exception of DID where concentrations were varied to maintain a consistent injection volume across all groups. A dose of 200mg/kg was used in all experiments with the exception of IEA experiment two (100 mg/kg) and DID studies (dose-response analysis ranging from 0-100 mg/kg). Tideglusib was suspended into vehicle by vortexing followed by bath sonication at 40 degrees Celsius for 60 minutes.

### Intermittent ethanol access, two-bottle choice

In experiment one, male and female mice (n=9-10/group) underwent two-bottle choice, intermittent ethanol access (IEA) for six weeks. In this model, mice always have access to two drinking bottles, one of which contains water. On Mondays, Wednesdays, and Fridays, one of the water bottles was replaced with 20% ethanol v/v at the beginning of the dark cycle and left on for 24 hours, with sides alternated each drinking session to avoid conditioning a side preference. Two hours (binge reading) and 24 hours after access to ethanol, volume measurements were taken, and ethanol consumption (g/kg), preference over water, and total fluid consumption (ml) were calculated. Baseline IEA consumption values (g/kg) were measured for three weeks. Mice were then acclimated to gavage treatments with three days saccharin administration followed by two days vehicle habituation before beginning gavage of 200mg/kg tideglusib or equivalent volume of vehicle on week six, after which effects on consumption and preference were measured for the week. Gavage treatments occurred 30-60 minutes prior to ethanol access.

In experiment two, to allow for greater power within groups, only male mice were included (n=12/group). Mice underwent the same two-bottle choice, IEA paradigm for a total of nine weeks. Baseline IEA consumption values (g/kg) were measured for three weeks. Mice were then acclimated to gavage treatments with two days saccharin administration followed by two days vehicle habituation before beginning gavage of 100mg/kg tideglusib or equivalent volume vehicle on week five, after which effects on consumption, preference, and total fluid intake were measured for four additional weeks. Two separate gavage treatments were administered each drinking day. The first treatment occurred two hours prior to ethanol access and the second occurred four hours following the first, occurring immediately after the two-hour binge read.

### Drinking in the dark

To establish a dose-response curve for tideglusib in a model of binge drinking behavior, male and female C57BL/6J mice (n=20/sex) were given access to 20% v/v ethanol for four hours as a single bottle access with no water present, four days a week, beginning three hours after lights off from Monday-Thursday. At the conclusion of the four-hour test, ethanol was removed and replaced with water bottles until the next experimental day. Baseline DID consumption values (g/kg/4hr) were measured for two weeks. Mice were then acclimated to i.p. injections with two days saline and four days corn oil before beginning injections of tideglusib.

For the first cycle of experimental drinking days, mice were randomly assigned to doses of tideglusib, including 0 mg/kg (corn oil vehicle), 10 mg/kg, 30 mg/kg and 100 mg/kg. One hour prior to ethanol access on experimental days, mice were weighed and injected with respective doses of tideglusib via i.p. injection. Animals then underwent a week of drug washout to confirm ethanol consumption returned to baseline values before rerandomizing groups and beginning a second cycle utilizing doses of 0 mg/kg, 50 mg/kg, 75 mg/kg and 100 mg/kg.

To assess time until washout of drug effects, a separate set of animals (n=24, males) underwent DID as described above. Mice received 100mg/kg tideglusib or corn oil vehicle via i.p. injection one hour prior to ethanol access (n=12/group) for one week, after which, mice then had an additional week of continued access to DID EtOH without any further tideglusib injections. Additionally, another group of mice (n=10/group, males) underwent this same DID testing except with a single bottle access to water during the testing period to assess if effects of tideglusib on consumption were specific to ethanol or fluid consumption in general.

### Liver function assessment

Blood serum was collected from male mice (n=3-5/group) with a history of nine weeks of water or ethanol DID. During the last week of DID, animals received i.p. injections of 100 mg/kg tideglusib or corn oil vehicle, and 24 hours after their last injection, blood was collected retro-orbitally. Serum was isolated and assessed for a panel of liver enzymes as a measure of liver functions, including alanine aminotransferase (U/L), aspartate aminotransferase (U/L), and alkaline phosphatase (U/L), using Vetscan preventative care profile plus kits (Zoetis US, Parsipanny, NJ).

### Blood ethanol content

Male mice (n=25) were gavaged with tideglusib or 26% peg-400 (Sigma), 15% Cremophor EL (Sigma), and water vehicle and 30 minutes later i.p. injected with 2.0 g/kg ethanol. Blood was collected via submandibular cheek punch at 10, 30, 60, and 90-minute time points post-ethanol (n=3-4/treatment/time). Blood was collected in BD microtainer tubes containing EDTA (Fisher Scientific, Waltham, MA) to prevent clotting and stored at −20 degrees Celsius until analysis. BECs were assessed by the VCU Analytics Core using headspace gas chromatography(23). EtOH content was calculated based on normalization to a consistent internal standard of 1-proponal in each sample and reported as mg/L.

### Taste Preference

Male and female mice (n=7-8/group) were gavaged with 200mg/kg tideglusib or vehicle for six days. They were given access to two bottles containing water and either saccharin or quinine one hour after tideglusib administration. Saccharin and quinine concentrations were 0.6mM and 25uM respectively for days 1-3 and were increased to 1.2mM and 40um for days 4-6. Quinine or saccharin preference compared to water was measured daily.

### Light/Dark Box

Two to three weeks prior to undergoing taste preference testing, the same male and female mice (n=4-8/group) were gavaged with 200mg/kg tideglusib or corn oil and 30 minutes later i.p. injected with 1.8 g/kg 20% v/v ethanol in saline or equivalent volume saline vehicle and put into a cage by themselves. Five minutes post-injection, mice were placed into the light side of light/dark boxes (Med Associates, Fairfax, VT) facing the dark compartment and assayed for 10 minutes. Percent time in light, percent distance traveled in light, and total locomotion were measured. The duration of the test was separated into two consecutive five-minute time bins.

### RNA sequencing analysis

A subset of mice from the second IEA experiment (n=4/group) were euthanized 24 hours after their last ethanol access to compare ethanol-drinking, tideglusib-treated mice to ethanol-drinking vehicle-treated mice. PFC was collected and sent for sequencing at the VCU Genomics Core. Following sequencing, samples were put through our pipeline to determine counts for each gene (*supplemental methods)*.

The paired end counts generated in the previous step were analyzed for differential expression analysis DESeq2(24). Genes with median counts of less than one across all samples were filtered out of the data and significantly differently expressed genes (DEGs) were determined using an uncorrected p<0.05 and a log-fold change (LFC) of LFC>0.1 or LFC<-0.1. Significant DEGs were then run through gene ontology analysis using ToppGene(25).

### Statistical Analysis

All statistical tests were performed in R. A p-value of 0.05 or lower was considered statistically significant.

## RESULTS

### A single dose of 200mg/kg tideglusib decreases ethanol consumption during two-bottle choice, IEA binge drinking

Prior to tideglusib administration, mice escalated their drinking during the three weeks of baseline, with no difference between groups. Following tideglusib administration, tideglusib-treated mice consumed significantly less ethanol than their vehicle-treated counterparts. Initial analysis consisted of a three-way repeated measures (RM) ANOVA with sex, treatment day, and treatment group as variables, and there was a main effect of sex on ethanol consumption (p=0.0001). This result was expected, as it is well established female mice statistically consume more ethanol than males(26–30). As a result, sexes were separated, and two-way RM ANOVAs were run on each separately.

Two-way RM ANOVA revealed a main effect of treatment day (p<0.0001) and a treatment group*treatment day interaction (p=0.007) in male mice on binge consumption (*Fig. 1a*) and a main effect of treatment day (p<0.0001) on binge ethanol preference (*Fig 1c*). Bonferroni post hoc analysis yielded significant differences in binge consumption on days 17-19 and binge preference on day 18 in tideglusib verse vehicle mice (p<0.05)

**Figure 1.**
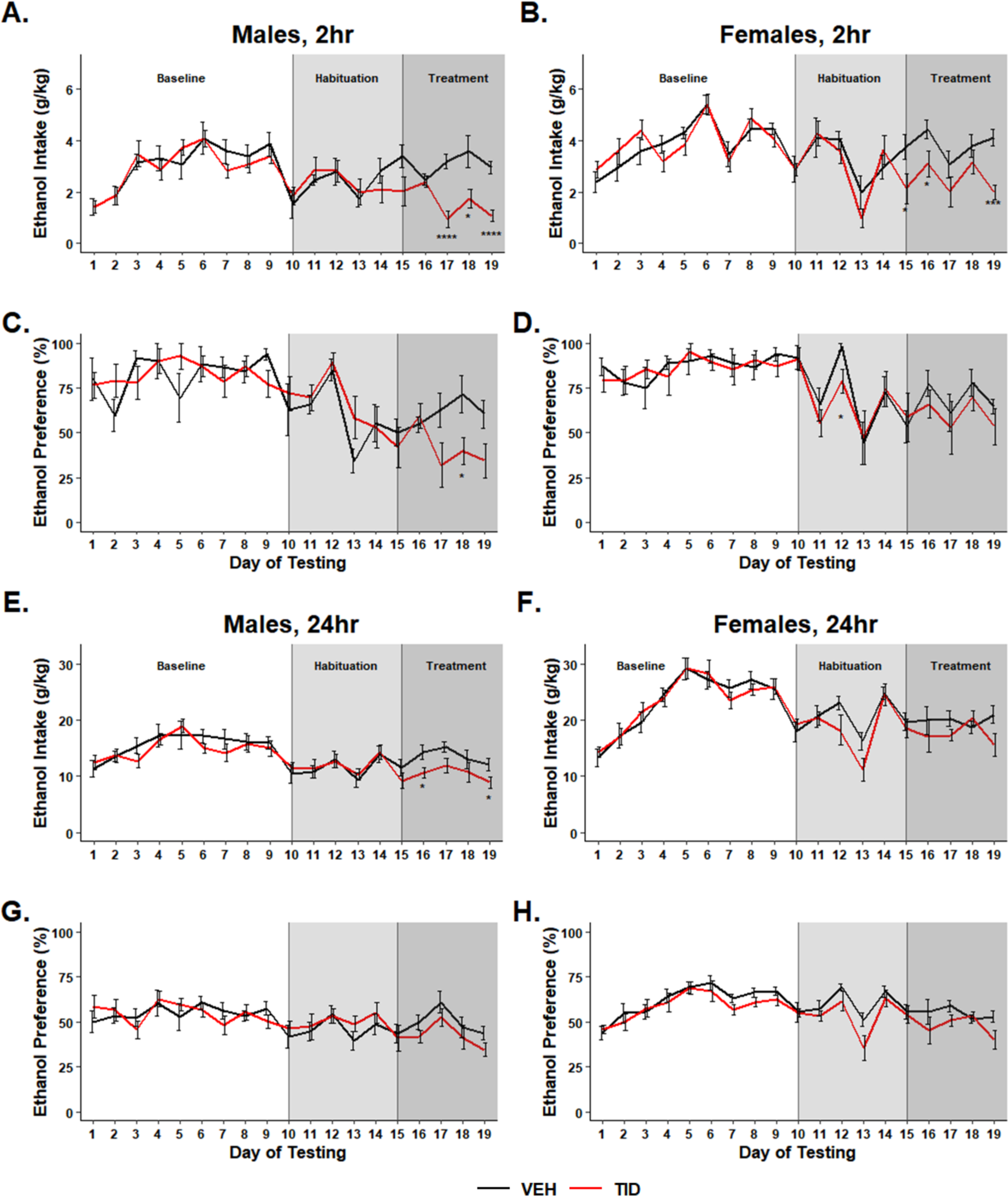
A single 200 mg/kg dose of tideglusib decreased binge ethanol consumption. Compared to vehicle-treated mice, tideglusib-treated mice (n=9-10/group) showed **A** a main effect of treatment day on decreased binge ethanol consumption (p<0.0001) and a treatment group*treatment day interaction (p=0.007) in males and **B** a main effect of treatment day on decreased binge ethanol consumption (p<0.0001) and a treatment-group*treatment-day interaction (p=0.02) in females. Bonferroni post hoc analysis showed significant differences between treatment groups on individual days (*p<0.05, ***p<0.001, ****p<0.0001). **C, D** There was a main effect of treatment-day on binge ethanol preference in both males and females (p<0.0001) but no effect of tideglusib. **E-H** There was an effect of treatment day on daily consumption and preference in both males and females (p<0.0001) but no effect of tideglusib treatment.

Female mice showed a main effect of treatment day (p<0.0001) and a treatment group*treatment day interaction (p=0.02) on binge consumption (*Fig. 1b*) and a main effect of treatment day (p<0.0001) on binge preference (*Fig. 1d*). Bonferroni post hoc analysis highlighted significant differences in binge consumption on days 15-16 and 19 and binge preference on day 12 in tideglusib verse vehicle mice (p<0.05)

This tideglusib dosing paradigm failed to elicit a main effect or interaction of tideglusib on ethanol drinking behaviors over the daily 24-hour ethanol access period, however, there was a main effect of day (p<0.0001). Bonferroni post hoc analysis revealed significant differences in male daily consumption on days 16 and 19 (p>0.05) but no differences in females (*Fig. 1e-h*).

### Two 100mg/kg doses of tideglusib decreases ethanol consumption and preference during two-bottle choice, IEA binge, and daily drinking

Prior to tideglusib administration, mice escalated their drinking during the four weeks of baseline, with no difference between groups. For the two-hour binge reading, two-way RM ANOVA showed a main effect of treatment group (p=0.029), treatment day (p<0.001), and treatment group*treatment day interaction (p<0.001) on consumption (*Fig. 2a*) and a main effect of treatment group (p=0.003), treatment day (p<0.001), and treatment group*treatment day interaction (p<0.001) on preference (*Fig. 2c*). Bonferroni post hoc analysis revealed significant differences between treatment groups during day three of baseline drinking and days 15-16, 18, and 20-24 during tideglusib treatment (p<0.05) for consumption and day three, 15-16, 19-24 for preference (p<0.05).

**Figure 2.**
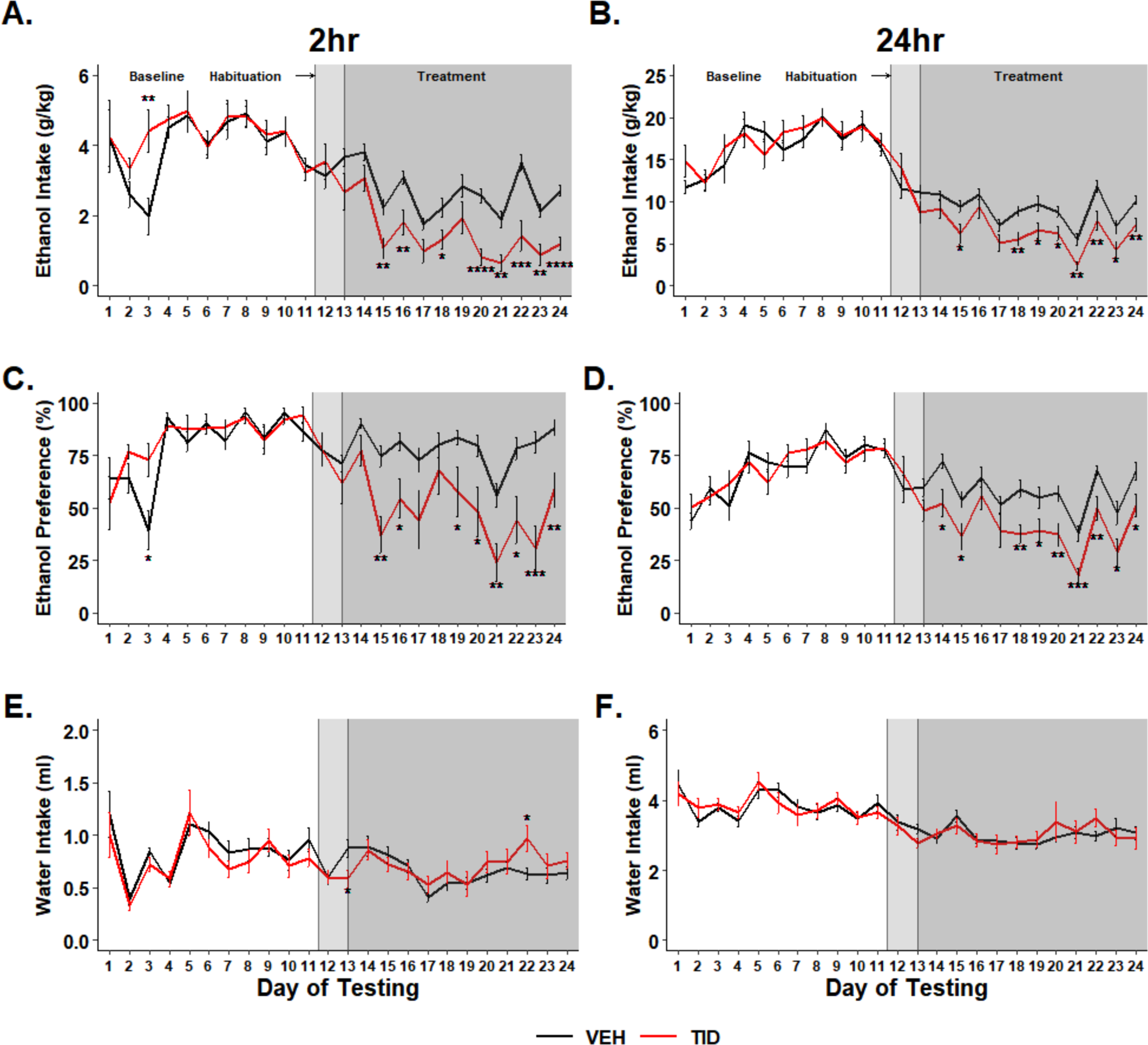
Two separate 100 mg/kg doses of tideglusib decreased binge and daily ethanol consumption and preference in males. Compared to vehicle-treated mice, tideglusib-treated mice (n=12/group, males) showed **A** a main effect of tideglusib treatment (p=0.029), an effect of treatment day (p<0.001), and a treatment group*treatment day interaction (p<0.001) on binge ethanol consumption and **B** an effect of tideglusib treatment (p=0.003), treatment day (p<0.001), and a treatment group*treatment day interaction (p=0.002) on daily ethanol consumption. **C** There was a main effect of treatment group (p=0.003), treatment day (p<0.001), and treatment group*treatment day interaction (p<0.001) in binge ethanol preference and **D** an effect of treatment group (p=0.032), treatment day (p<0.001) and treatment group*treatment day interaction (p<0.001) on daily ethanol preference. Bonferroni post hoc analysis showed significant differences between treatment groups on individual days (*p<0.05, ** p<0.01, ***p<0.001, ****p<0.0001). **E,F** There was a main effect of treatment day on water intake in both the binge and daily timepoints (p<0.05) but no effect of tideglusib treatment.

Tideglusib also significantly reduced daily ethanol behaviors under this dosing paradigm, with two-way RM ANOVA revealing an effect of treatment day (p<0.001) and a significant treatment group*treatment day interaction (p=0.002) on consumption (*Fig. 2b*), and an effect of treatment group (p=0.032), treatment day (p<0.001) and treatment group*treatment day interaction (p<0.001) on preference (*Fig. 2b*). Bonferroni post hoc analysis yielded significant differences between treatment groups during day three of baseline drinking and days 15 and 18-24 (p<0.05) for consumption and days 14-15 and 18-24 during tideglusib treatment for preference (p<0.05).

To assess effects of tideglusib on water consumption, an additional n=12/group mice were included who remained ethanol naïve throughout the course of the experiment and instead received two bottles of water to drink from. There was a main effect of treatment day on water intake in both the binge and daily timepoints (p<0.05) but no effect of tideglusib treatment (*Fig. 2e-f*), suggesting the actions of tideglusib are specific to decreasing ethanol intake and is not likely to cause issues with overall fluid intake in our model.

### Tideglusib treatment decreases ethanol consumption during DID

There was an effect of sex during DID (p<0.0001) where females consumed more g/kg EtOH than males, as well as an effect of tideglusib dose (p<0.0001) according to two-way ANOVA. Bonferroni post hoc analysis revealed a dose of 50 mg/kg (p=0.017), 75 mg/kg (p<0.0001), or 100 mg/kg (p<0.0001) was sufficient in decreasing consumption in male mice compared to a vehicle control. There was no further decrease in consumption with 100 mg/kg compared to 75 mg/kg. In females, a dose of 75 mg/kg (p=0.0002) or 100 mg/kg (p<0.0001) significantly decreased consumption compared to vehicle, however, the higher dose of 100 mg/kg was significantly different than the 75 mg/kg dose (p=0.009) (*Fig. 3a*).

**Figure 3.**
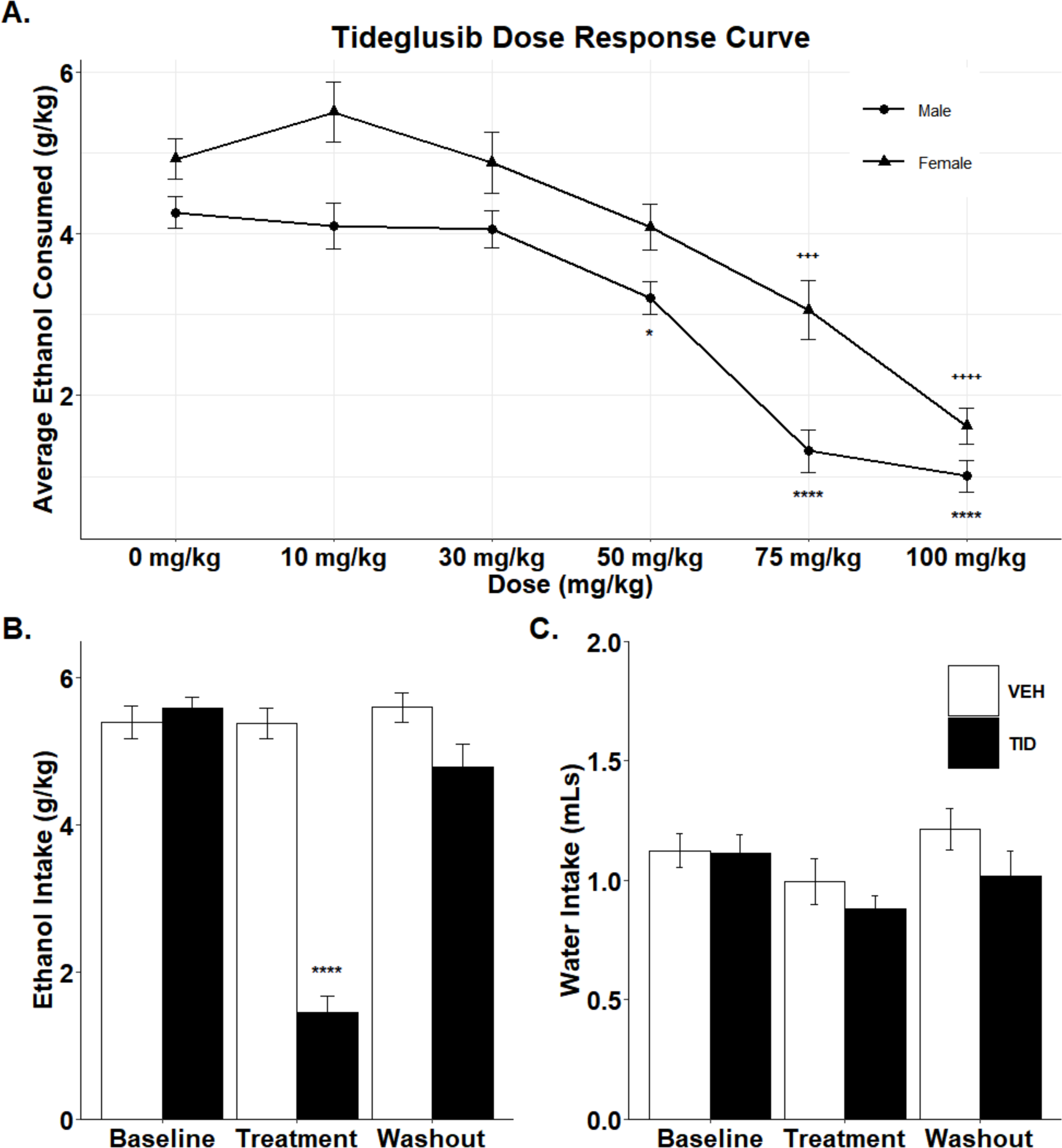
Tideglusib decreased ethanol consumption during drinking-in-the-dark without affecting water intake. **A** Tideglusib treatment decreased ethanol consumption in a dose-dependent manner (p<0.0001). Females consumed more g/kg EtOH than males over the course of the study (p<0.0001) (n=10-20/sex/dose). * denotes significance between dose and vehicle control in males and + denotes significance in females (* p<0.05, +++p<0.001, ****/++++p<0.0001). **B** 100 mg/kg of tideglusib decreases ethanol consumption during the treatment phase compared to baseline consumption and drug washout. Tideglusib treatment is also significantly different from vehicle treatment during the treatment period (****p<0.0001). **C** Tideglusib treatment has no effect on water intake.

Using the most effective dose from both sexes (100 mg/kg), two-way ANOVA revealed tideglusib significantly decreased ethanol consumption compared to baseline values, and tideglusib-treated mice drank significantly less than vehicle-treated mice during the treatment period (p < 0.0001). Prior to tideglusib treatment, mice assigned to either treatment group consumed equal amounts of ethanol in DID. Following washout, tideglusib-treated mice returned to baseline consumption values, and there was no significant difference between drug versus vehicle-treated animals during washout (*Fig. 3b*). Tideglusib had no effect on water intake (*Fig. 3c*).

### Tideglusib treatment has minimal effects outside ethanol consumption

Two-way ANOVA showed there was no effect of tideglusib treatment on levels of alanine aminotransferase (*Supplemental Fig. 1b)*, or aspartate aminotransferase (*Supplemental Fig. 1c)*. However, tideglusib did decrease levels of alkaline phosphatase (p=0.0008) (*Supplemental Fig. 1d)*. Tukey’s HSD post hoc revealed a significant difference between water-drinking animals treated with tideglusib versus vehicle (p=0.0478) and between ethanol-drinking tideglusib versus vehicle (p=0.0249).

There was a significant effect of time following ethanol injection on blood ethanol concentrations (BECs) according to two-way ANOVA (p<0.0001), where later timepoints had lower BECs, as to be expected. There was no effect of tideglusib treatment compared to vehicle controls nor a treatment*time interaction, suggesting tideglusib does not alter ethanol pharmacokinetics (*Supplemental Fig. 1a)*. Tideglusib also did not alter taste preference for quinine nor saccharin at either the low or high concentrations tested according to two-way ANOVAs run for each sex and tastant separately (*Fig. 4a-b*).

**Figure 4.**
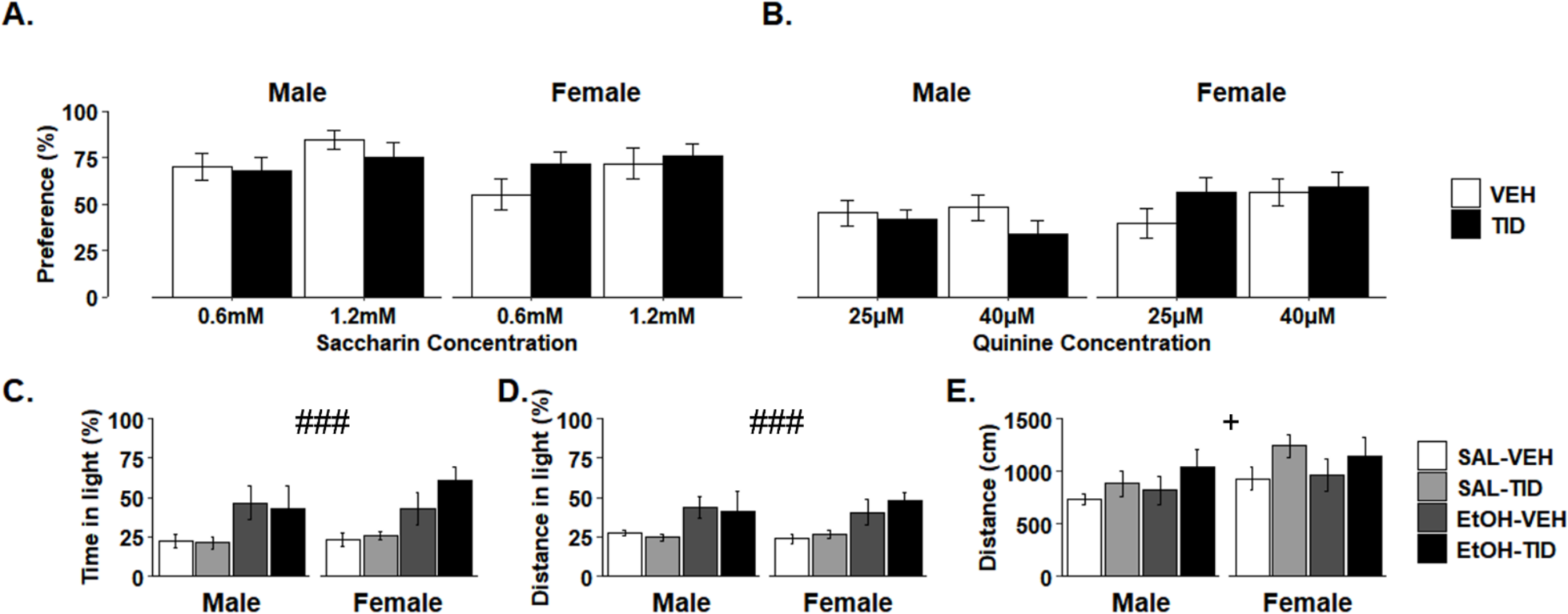
Tideglusib has no effect on taste preference or anxiety-like behavior but transiently increases total locomotion. **A** Tideglusib treatment does not change taste preference in either sex (n=7-8/group) for saccharin at 0.6mM or 1.2mM **B** nor quinine at 25µM or 40µM concentrations. **C,D** i.p. ethanol significantly decreases anxiety-like behavior in the light-dark box (###p<0.001) with no effect of sex, but there is no effect of tideglusib (n=4-8/group). **E** Tideglusib transiently increases total locomotion in the light-dark box (+p=0.46) however, no comparisons reached significance in post hoc testing.

In the light/dark box, there was a main effect of ethanol on anxiety-like behaviors as measured by a three-way ANOVA (p<0.001), but tideglusib treatment did not alter anxiety-like behavior as measured by either percent time (*Fig. 4c*) or percent distance in light (*Fig. 4d*), nor was there a treatment*ethanol interaction. Tideglusib did significantly increase locomotion during the first 5 minutes of the assay (p=0.047) (*Fig. 4e*), but there was no effect during the last 5 minutes of the assay, nor when the entire 10-minute test was compared (data not shown). There was no effect of sex on any measure.

### Tideglusib treatment regulates transcription factor binding sites involved in the Wnt signaling pathway

Differential expression analysis revealed 711 DEGs in the PFC between ethanol drinking animals treated with tideglusib versus vehicle (*supplemental table 1*). Gene ontology analysis in ToppGene revealed significant functions of these genes in multiple transcription factor binding sites (TFBS) (*supplemental table 2*). The top five TFBS all showed an overrepresentation of genes involved in Wnt signaling (*Fig. 5a*). In particular, the two most significant TFBS, transcription factor three (TCF3) and lymphoid enhancer-binding factor (LEF1), are both binding sites for β-catenin, a known member of the canonical Wnt-signaling pathway along with GSK3β (31). There was a total of 119 significant DEGs between ontology results for LEF1 and TCF3, with 22 of these genes being shared between the two. The significant genes (p<0.05) with the ten greatest log(2)-fold changes contributing to the ontology results of TCF3 and LEF1 were identified and are presented in figure 5b.

**Figure 5.**
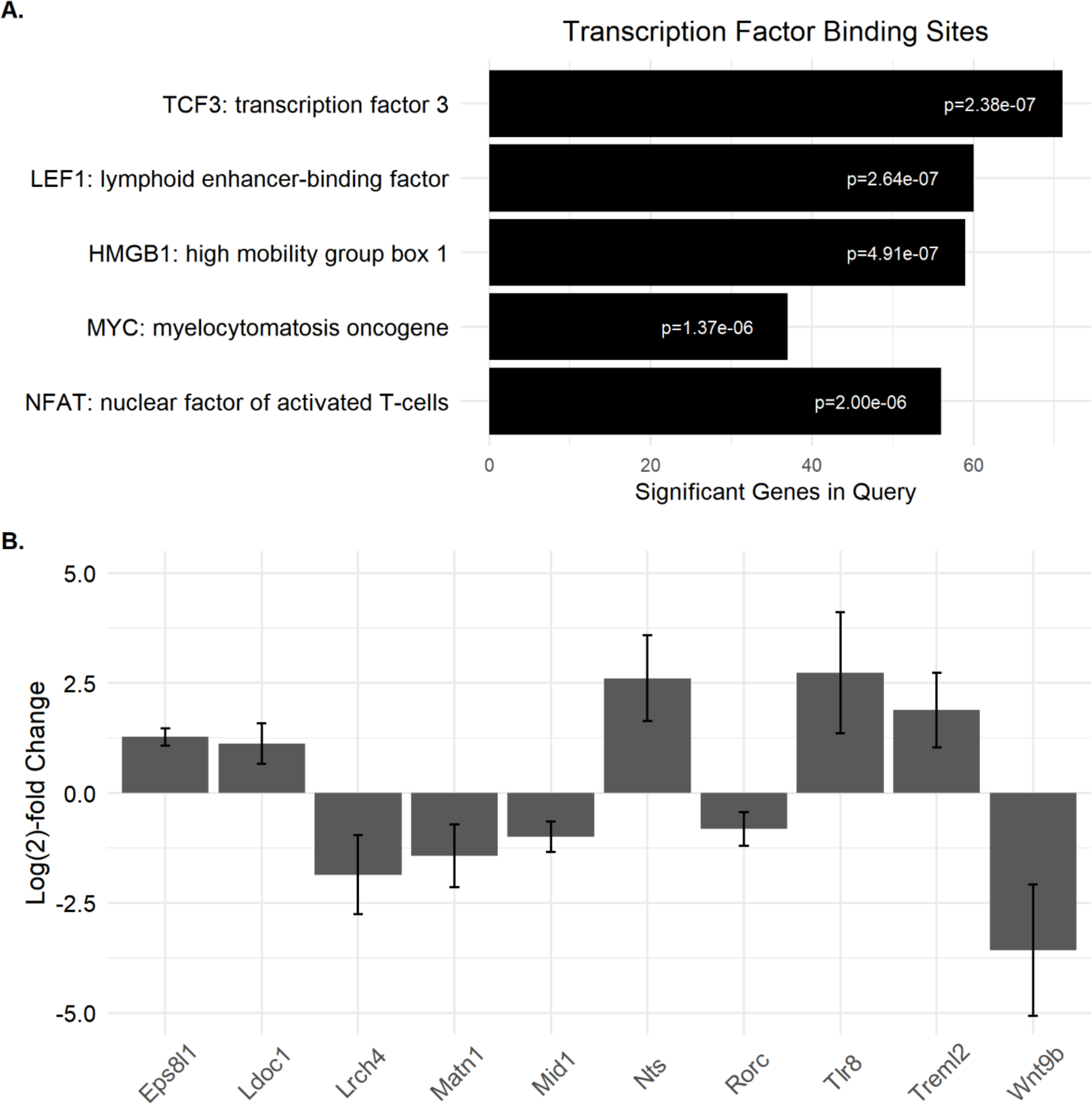
Tideglusib-treated mice show differential gene expression of genes related to β-catenin binding. **A** The top five gene ontologies for transcription factor binding sites in ethanol-drinking mice treated with tideglusib vs vehicle (n=4/group) and their significant p-values are shown. All five are involved in Wnt signaling, with TCF3 and LEF1 binding β-catenin as part of the canonical Wnt signaling pathway. **B** The top ten differentially expressed genes contributing to the ontology results of TCF3 and LEF1 as determined by greatest log(2)-fold change (all p>0.05) were plotted. The log(2)-fold change represents differences in expression within tideglusib-treated animals vs vehicle.

## DISCUSSION

GSK3β has been linked to multiple processes involved in neurotransmission, including synapse formation and plasticity(10). Additionally, work from our lab has previously shown modulations in GSK3β abundance or activity modulate ethanol behaviors(6). In the present study, we showed GSK3β inhibition via tideglusib significantly decreases ethanol consumption in two different drinking models: the IEA model of progressive alcohol consumption and the DID model of binge drinking. Importantly, tideglusib had no effects on taste preference, anxiety-like behavior, or ethanol metabolism, though there was a transient effect on locomotion.

Previous work shows inhibition of GSK3β via drugs such as lithium and TDZD-8 successfully decrease ethanol consumption(6). While lithium is commonly used in the clinic for treatment of bipolar disorder, it is not selective for GSK3β alone, also targeting GSK3α, phosphoinositide hydrolysis, adenylyl cyclase, G protein, and protein kinase C(32). This promiscuity could be the reason for the high occurrence of negative side effects while on lithium. Lithium also has a high rate of nonadherence in patients(33, 34). Conversely, TDZD-8 is highly selective for GSK3β, however, it is not currently clinically available and has poor bioavailability.

Tideglusib is both selective for GSK3β and clinically available. Phase 2 clinical trials for treatment of Alzheimer’s disorder, progressive supranuclear palsy, autism spectrum disorder, and myotonic dystrophy 1 have been completed, and phase 2/3 trials for congenital myotonic dystrophy are currently underway(18–22). The most frequent side effects in these trails were diarrhea in 13-18% and an asymptomatic, transient, and reversible increase in aminotransferase in 9-16% of subjects(35–37). In our studies, we assessed levels of alanine aminotransferase, aspartate aminotransferase, and alkaline phosphatase. There was no effect of tideglusib treatment on either aminotransferase, however, we did see a tideglusib-induced increase in alkaline phosphatase (*Supplemental Fig. 1b-d*). A clinical trial on myotonic dystrophy also reported mild nasopharyngitis in 31% of patients(37), though this effect was not observed in trials on other disorders. The findings in all trials are consistent; tideglusib is safe and well tolerated within human populations, a pivotal aspect in furthering the study of tideglusib in the use of AUD.

Here we have shown tideglusib successfully decreases both ethanol consumption and preference in mice habituated to drinking alcohol within the IEA model. IEA is well documented to result in progressive, increased consumption over time(38). This behavior is often seen in AUD, where initial exposure to alcohol progresses to levels which results in excessive and compulsive consumption. This suggests tideglusib may have utility in decreasing drinking behaviors in humans with AUD. Our first IEA experiment gave mice a single 200mg/kg dose of tideglusib. This elicited a decrease in drinking behaviors during the binge consumption period (*Fig. 1a-d*) but failed to have a significant effect on daily consumption or ethanol preference (*Fig. 1e-h*). This is likely due to the relatively short half-life of tideglusib. While these studies were being conducted, a report determined the half-life for tideglusib in male Balb/C mice was 5.12 hours(39). We verified this in C57BL/6J mice (4.9 h, data not shown), and therefore we performed a second IEA experiment where we administered two separate 100mg/kg doses of tideglusib spread four hours apart to elicit changes in consumption over a 24hr period. This dosing paradigm produced significant decreases in both binge and daily ethanol consumption and preference (*Fig. 2*).

Having shown tideglusib reduces ethanol behaviors in a progressive model of AUD, we next moved to the DID model of binge drinking. In this model mice consume high levels of ethanol within a relatively short time span, mirroring human binge drinking behavior(40). Our study showed tideglusib was fast acting to decrease consumption in mice habituated to ethanol drinking. Following washout of tideglusib, mice returned to baseline levels of consumption (*Fig. 3*). This shows not only is tideglusib rapid to enact its effects on drinking, but it is also equally rapid to cease these actions without causing long term modulations in drinking behavior. This suggests GSK3β inhibition via tideglusib may have utility as an intervention to lower ethanol consumption immediately prior to onset of a return to alcohol in patient populations recovering from AUD.

Ethanol metabolism is related to the amount of ethanol consumed, so to assess whether changes in ethanol consumption were due to change in metabolism, we assessed BECs following a high-dose ethanol injection. Here we have shown GSK3β modulation via tideglusib has no effects on metabolism as measured via BEC time course (*Supplemental Fig. 1a*).

To further assess the potential of tideglusib as a therapeutic agent, we measured effects of global GSK3β inhibition on behaviors outside of ethanol-consumption. Certain behavioral phenotypes could potentially alter ethanol behaviors without necessarily being involved in ethanol consumption. For this reason, we measured effects of tideglusib on taste preference, anxiety-like behavior, and locomotion.

An animal’s alcohol consumption can be altered by an animal’s taste preference, as perceived taste of ethanol is an important determinant of consumption(41). We measured taste preference for saccharin, a sweet substance which mice voluntarily consume under baseline conditions, and quinine, a bitter substance which elicits decreased self-administration. Saccharin was selected over sucrose to avoid confounds of increased caloric load from sucrose. There was no change in taste preference for either adulterant (*Fig. 4b,c*), suggesting tideglusib is likely not reducing alcohol consumption by altering taste in the mouse.

It is well established there is a high prevalence of comorbidity between alcohol use disorders and anxiety disorders(42, 43). One hypothesis as to the development of this etiology is the “tension-reduction” hypothesis, which states alcohol produces a reduction in anxiety symptoms, and this reduction is a main motivator for drinking behavior(42, 44). Previous studies which decreased GSK3β levels using transgenic mouse lines found reduced anxiety-like responses(45, 46). Similarly, studies in rat nucleus accumbens shell show GSK3β knockdown decreased sucrose neophobia and cold stress defecation, suggesting possible effects on anxiety(47).

We therefore measured effects of global GSK3β inhibition on basal anxiety, as changes in anxiety could potentially account for the changes we see in consumption following tideglusib. In our studies, we demonstrated the anxiolytic effects of ethanol, but importantly, there were no effects of tideglusib treatment on anxiety-like behavior nor any interaction with tideglusib and alcohol (*Fig. 4d,e*). The systemic nature of our GSK3β modulations compared to the more targeted approaches used in the studies mentioned above may account for the differences we found regarding effects on anxiety as studies have shown GSK3β modulations to have differential effects depending on brain region and cell type specificities(46, 48, 49).

Locomotion is another important behavior to measure as effects in movement can affect ability to obtain alcohol. Tideglusib did produce a transient increase in total distance traveled within the first five minutes of testing (*Fig. 4f*). However, this effect on locomotion was quick to dissipate, with no effect on the last five minutes of testing or when the entire ten-minute test was viewed as a whole (data not shown). This suggests the locomotor increase may not be clinically relevant. Other preclinical studies on GSK3β modulations via genetic knockdown or pharmacological inhibition have shown no effects on locomotion(45, 50) and similarly clinical trials in humans have not reported tideglusib-induced movement effects in patients(35–37). It should be noted in a clinical trial for progressive supranuclear palsy, the patient population displays movement abnormalities pre-treatment. However, while tideglusib was ineffective at improving motor function in these subjects, no new or worsening effects on movement were reported(35).

Tideglusib-induced inhibition of GSK3β may be eliciting its actions on ethanol consumption via effects on the Wnt signaling, particularly through the canonical Wnt-β-catenin signaling pathway. When GSK3β is active, it phosphorylates β-catenin and β-catenin is sequestered in a destruction complex with GSK3β, AXIN 1/2, adenomatous polyposis coli (APC), and casein kinase I-alpha (CK1α), where it is targeted for ubiquitylation. When GSK3β is inhibited, β-catenin is no longer targeted for degradation, allowing it to enter the nucleus and carry out its transcription factor activities by binding to the TCF/LEF transcription factor family(31, 51–55). Wnt-β-catenin signaling is implicated in cell proliferation across many developmental stages, including in adult tissue and has become an interesting target in the study of plasticity(56, 57). Wnt signaling via Wnt-3a and Wnt-7a have both been shown to regulate presynaptic plasticity via the canonical Wnt-β-catenin signaling pathway by strengthening excitatory synapses(58–60). Interestingly, lithium has been shown to activate Wnt signaling(50) and induce the same changes in presynaptic remodeling we see with Wnt-7a signaling(61). Additionally, both Wnt-3 and Wnt-7a have been shown to enhance long-term potentiation post-synaptically as well(62, 63), and treatment with Wnt-3a-induced transcriptional changes in genes associated with learning and memory and neurotransmitter release(64). Altogether the Wnt-β-catenin signaling pathway seems to be a target in regulating neuronal plasticity, and GSK3β inhibition may be preventing alcohol-induced changes to canonical Wnt signaling.

Alcohol use disorder affects more than 29.5 million people within the United States alone according to DSM-5 criteria(1). Despite the levels of AUD within the population, only 7.6% of individuals with AUD received any treatment in 2022, and only 2.1% of people with AUD received medication-assisted treatment(1). A part of this treatment gap can be attributed to the lack of effective therapeutic medications. Currently, there are three FDA-approved medications for the treatment of AUD (disulfiram, naltrexone, and acamprosate), and these medications are not very effective. There is need for a new, more effective therapeutic for AUD treatment, and the data presented here support the use of tideglusib as such.

## ACKNOWLEDGEMENTS

We thank the laboratory of Dr. Arun Sanyal, in particular members Mulugeta Seneshaw and Faridoddin Mirshahi, for their guidance on liver enzyme analysis.

## AUTHOR CONTRIBUTIONS

Conceived and designed the study: SG AVDV JTW MFM. Acquired the data: SG AVDV AH DB AM BO. Analyzed the data: SG WR MFM. Wrote the paper: SG MFM.

## Supplemental Methods

### RNA isolation and sequencing

Mice were euthanized 24 hours after their last ethanol access via cervical dislocation and decapitation. PFC was collected, flash-frozen in liquid nitrogen, and stored at −80° C. RNA was extracted and purified using RNeasy mRNA kits (Qiagen, Germantown, MD). Quality of isolated RNA was assessed using both a NanoDrop assay (Thermo Fisher Scientific, Waltham, MA) and BioAnalyzer 2100 (Agilent Technologies, Santa Clara, CA). Isolated RNA samples were then shipped on dry ice for poly-A selected library construction and sequencing on an Illumina NextSeq 2000 platform with P3 flow cell (Illumina, Inc. San Diego, California) in a strand-specific 100bp paired-end sequencing reaction at the VCU Genomics Core.

Default read quality control, duplicate sequence identification, adapter sequence trimming, and G-C content settings were used for fastP processing (v0.23.2). Read strandedness was determined by default parameters in the how_are_we_stranded_here Python library (v1.0.1). Reads were aligned to GRCm39 genome by STAR (v2.7.10b). Sorting of BAM files and downstream indexing was performed with Samtools (v1.5.1). The featureCount program in the Subread package (v2.0.1) was used to quantify aligned reads. Samples were screened for outliers by principal component analysis and visualization of variance stabilized-transformed reads.

**Supplemental Figure 1.**
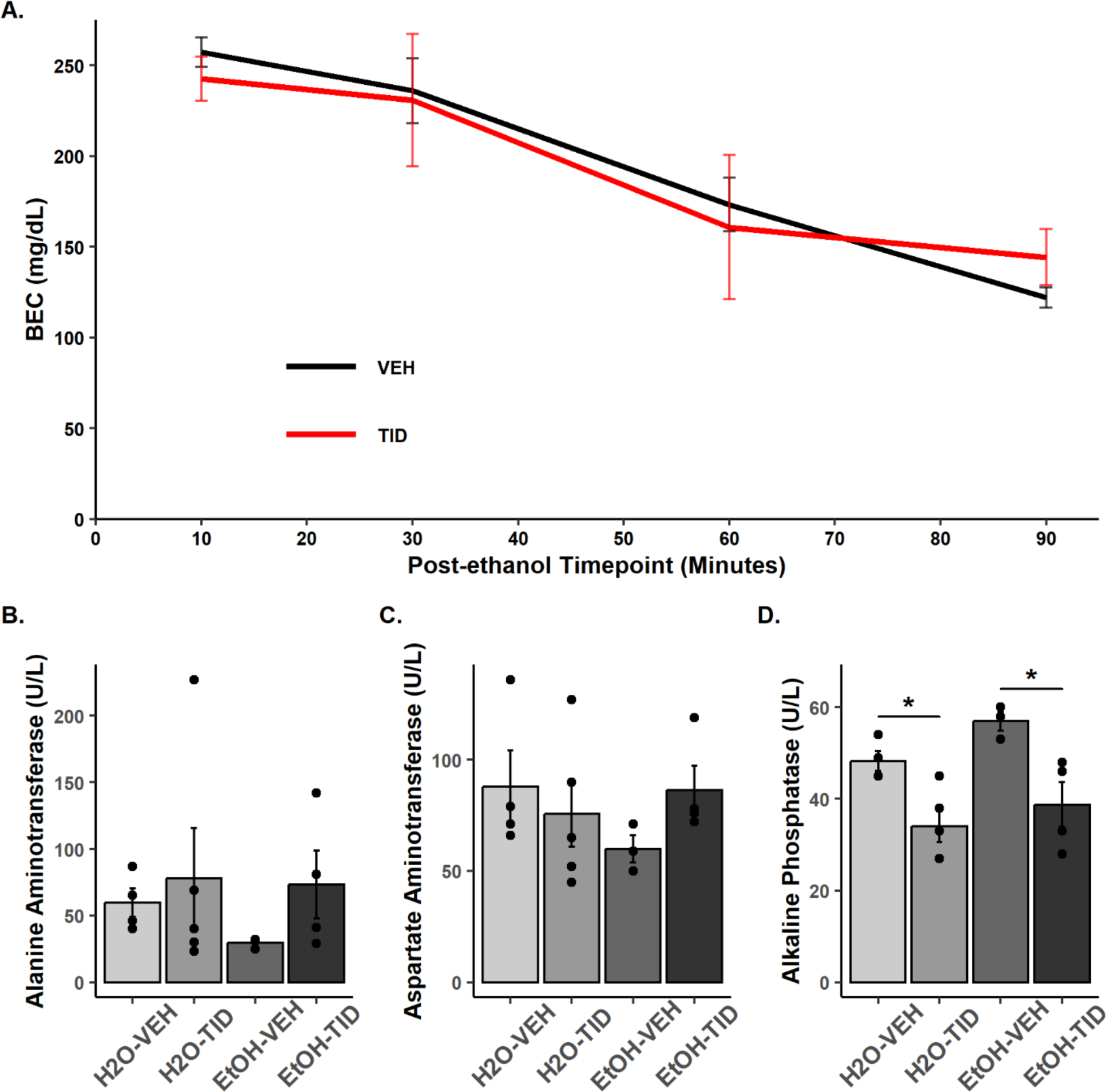
Tideglusib-treatment has no effect on ethanol metabolism or aminotransferase levels but increases alkaline phosphatase levels. **A** There was a main effect of time post injection on BEC (pp<0.0001) but no effect of tideglusib on ethanol pharmacokinetics as measured by BEC at any timepoint tested (10, 30, 60, 90min) (n=3-4/group/timepoint). **B,C** Tideglusib has no effect on alanine aminotransferase or aspartate aminotransferase but **B** significantly decreases alkaline phosphatase levels (p=0.0008) (n=3-4/group). There was no effect of ethanol on any enzyme measured nor an interaction between ethanol and tideglusib.

